# Emergence of a New ST617 Hypervirulent *Klebsiella pneumoniae* with ESBL^+^ and Strong Biofilm Forming Ability in China

**DOI:** 10.1101/2025.04.03.646950

**Authors:** Yuyun Yu, Xu Dong, Yanghui Xiang, Yi Li, Kefan Bi, Jiaying Liu, Lin Sun, Tiantian Wu, Ying Zhang

**Affiliations:** State Key Laboratory for Diagnosis and Treatment of Infectious Diseases, National Clinical Research Center for Infectious Diseases, Collaborative Innovation Center for Diagnosis and Treatment of Infectious Diseases, The First Affiliated Hospital, Zhejiang University School of Medicine, Hangzhou, China; Jinan Microecological Biomedicine Shandong Laboratory, Jinan, China; Jiangsu Key Laboratory of Zoonosis/Jiangsu Co-Innovation Center for Prevention and Control of Important Animal Infectious Diseases and Zoonoses, Yangzhou University, Yangzhou, China; Key Laboratory of Prevention and Control of Biological Hazard Factors (Animal Origin) for Agrifood Safety and Quality, Ministry of Agriculture of China, Yangzhou University, Yangzhou, China; Joint International Research Laboratory of Agriculture and Agri-Product Safety, Yangzhou University, Yangzhou, China

## Abstract

The increasing infections caused by hypervirulent *Klebsiella pneumoniae* (hvKP) resistant to carbapenem, broad-spectrum β-lactam and colistin has posed great challenges to global public health. However, the typing and detailed molecular features of the multi-drug resistant hvKP remain to be improved. Here, we characterized a broad-spectrum β-lactam-resistant *K. pneumoniae* ST617 isolate that has not been previously reported to cause severe disease in China. We obtained the whole-genome sequence of one *K. pneumoniae* ST617 isolate—HZ7—from a patient with bile duct infection. Phylogenetic reconstruction was performed to investigate the possible host evolutionary origin of HZ7, and the virulence property was studied in comparison with the nearest ST617 neighbours—YZ strains isolated from patient blood infections. Our studies indicated that the HZ7 is an ESBL^+^ Kp that is likely evolved from environmental or animal-derived sources. Importantly, the virulence assay showed that HZ7 was highly virulent and compared with the well known hvKP strain—NTUH-K2044 and the highly virulent clinical strain ST11-K64—HZ54—isolated from a patient with severe pneumonia. In addition, HZ7 showed strong biofilm formation ability. Whole-genome sequencing analysis identified seven potential virulence genes in HZ7: kp7_000748, kp7_002156, kp7_002157, kp7_002158, kp7_002159, kp7_003103, kp7_003104, which are absent in *K. pneumoniae* patient isolates from Yangzhou and the environmental samples. Finally, transcriptomic analysis of HZ7 identified six genes—*wcaJ, fimA, mrkH, pgaA, ugd*, and *gndA* that are associated with biofilm formation. Given its high virulence likely associated with the seven unique genes acquired via horizontal transfer and strong biofilm forming ability, this strain should be closely monitored to prevent wider spread in the clinic.

## Introduction

*Klebsiella pneumoniae* is an opportunistic pathogen that exists in immunocompromised populations and is an important factor in community-acquired infections as well as a major cause of nosocomial pneumonia[1]. In addition to pneumonia, it can cause a variety of infections, including meningitis, bacteremia, and urinary tract infection[2], and it is more common in diabetic patients and alcoholics[3]. In the past two decades, in addition to the classic *Klebsiella pneumoniae* (cKP), another new “highly virulent” *K. pneumoniae* (hvKP) infection rate has gradually appeared in the public eye[4]. It is worth noting that hvKP can cause invasive infections in young, healthy individuals, liver abscesses[5] or infections such as endophthalmitis and meningitis[6].

The typing of *K. pneumoniae* is based on multilocus sequence typing (MLST) of seven housekeeping genes (*gapA, infB, mdh, pgi, rpoB, phoE*, and *tonB*)[7]. *K. pneumoniae* ST11 is the most important clinical strain in China[8]. As the main virulence factor of *K. pneumoniae*, the capsular polysaccharide can be divided into 82 K serotypes. Serotypes K1 and K2 were found to be the most virulent[9]. ST11-K47 and ST11-K64 that are highly virulent and carbapenem-resistant *K. pneumoniae* have been repeatedly reported in bacterial liver abscesses[10-12].

Since the first case of hvKP was first detected in Taiwan in 1986[13], the incidence of hvKP in the Asia-Pacific region has gradually increased[14]. In recent years, carbapenem[15, 16], colistin[17], and broad-spectrum β-lactam [18] resistant highly virulent *K. pneumoniae* have been reported frequently. According to the recent annoucement from the World Health Organization, the emergence of multi-drug resistant high-virulence *K. pneumoniae* strains has brought significant challenges to treatment and pose a huge threat to global public health[19, 20].Therefore, it is critical to identify and characterize different types of antibiotic resistant hvKP.

In this study, we elucidate the genomic structure, virulence, and biofilm formation characteristics of an ESBL^+^-hvKP clinical strain at a tertiary hospital in the Zhejiang province of China. The mechanism of the high virulence and biofilm formation were analyzed based on whole-genome sequencing and RNA-seq analyses.

## Results and Discussion

### Clinical isolation of *K. pneumoniae* strains

Patient 1 (where *K. pneumoniae* HZ7 strain was isolated) was a 35-year-old woman, who had a 3-year history of liver cancer. She was admitted to hospital due to abdominal distension and yellow skin for 1 week. She had recurrent fevers while in hospital and was treated with indomethacin and celecoxib. One bacterial isolate was obtained from bile and was identified as *K. pneumoniae* ST617 strain. Meropenem was used for anti-infective therapy during hospitalization. The patient was discharged after 2 weeks, who had a poor prognosis. Strains YZ2429, YZ2487 and YZ22127 were all isolated from patient blood infections (Patient 2, 3, 4) in Yangzhou[21], and were all identified as serotype ST617 by sequencing. They were respectively admitted to hospital with multiple injuries, severe acute pancreatitis, and cerebral haemorrhage and were discharged after treatment. Patient 5 (strain HZ54) was a 63-year-old man, admitted to hospital with septic shock after a car accident. And blood cultures suggested CR-KP and was identified as *K. pneumoniae* ST11-KL64 strain. The virulence gene profiles of the 5 clinical strains were characterized using sequence alignment against the VFDB database. Blast analysis showed that HZ54 contains 3 known hvKP-related genes, including *rmpA2, ybt9*, and *iuc1*, but isolates HZ7, YZ2429, YZ2487 and YZ22127 did not contain these known virulence genes (Table S1).

### Characteristics of serotype ST617 strains

Strains YZ2429, YZ2487 and YZ22127 have been reported in previous studies as positive for *blaKPC-2* and were resistant to most commonly used antibiotics[20]. The MIC of HZ7 was determined by using broth microdilution method and was interpreted according to CLSI clinical breakpoints. HZ7 was resistant to Cotrimoxazole, Ceftazidime, Cefepime, Cefuroxime axetil, Cefuroxime, Cefazolin, Ceftriaxone, Aztreonam, and Ampicillin, but susceptible to Imipenem, Ertapenem, Cefoxitin, Gentamicin, Tigacycline, Ciprofloxacin, Amikacin, and Kanamycin. In particular, we found HZ7 was positive for extended-spectrum beta-lactamase (Table 1).

### Virulence study of *K. pneumoniae* strain HZ7 in comparison with other strains

It has been reported that *K. pneumoniae* ST11-KL64 is identified as hypervirulent *K. pneumoniae* (hvKP)[22, 23]. Hence, we selected a clinical strain (HZ54) with the ST11-KL64 serotype isolated from drainage fluid from a patient as a high virulence strain. NTUH-K2044(ST23/KL1) was selected as a typical representative of hvKP[17]. The environmental strain CGMCC1.839 from Donghu farm in Wuhan was used as a non-pathogenic strain for comparison. Galleria mellonella larvae have widely been used as an alternative model organism for bacterial pathogens, and the virulence of *K. pneumoniae* isolates was assessed in Galleria mellonella model. We compared survival rates of Galleria mellonella infected with HZ7, YZ2429, YZ2487, YZ22127, the high-virulence strains NTUH-K2044 and HZ54 and low-virulence strain CGMCC1.839 to evaluate their differences in pathogenicity. As shown in Figure 1(A), the larvae survival rates were much lower following infection with HZ7, HZ54 and NTUH-K2044, as compared to CGMCC1.839. The HZ7 strain was more virulent than the other three ST617 strains (YZ2429, YZ2487, YZ22127) with Galleria mellonella survival <40% at 24 h and ≤15% at 60 h.

**Figure 1.**
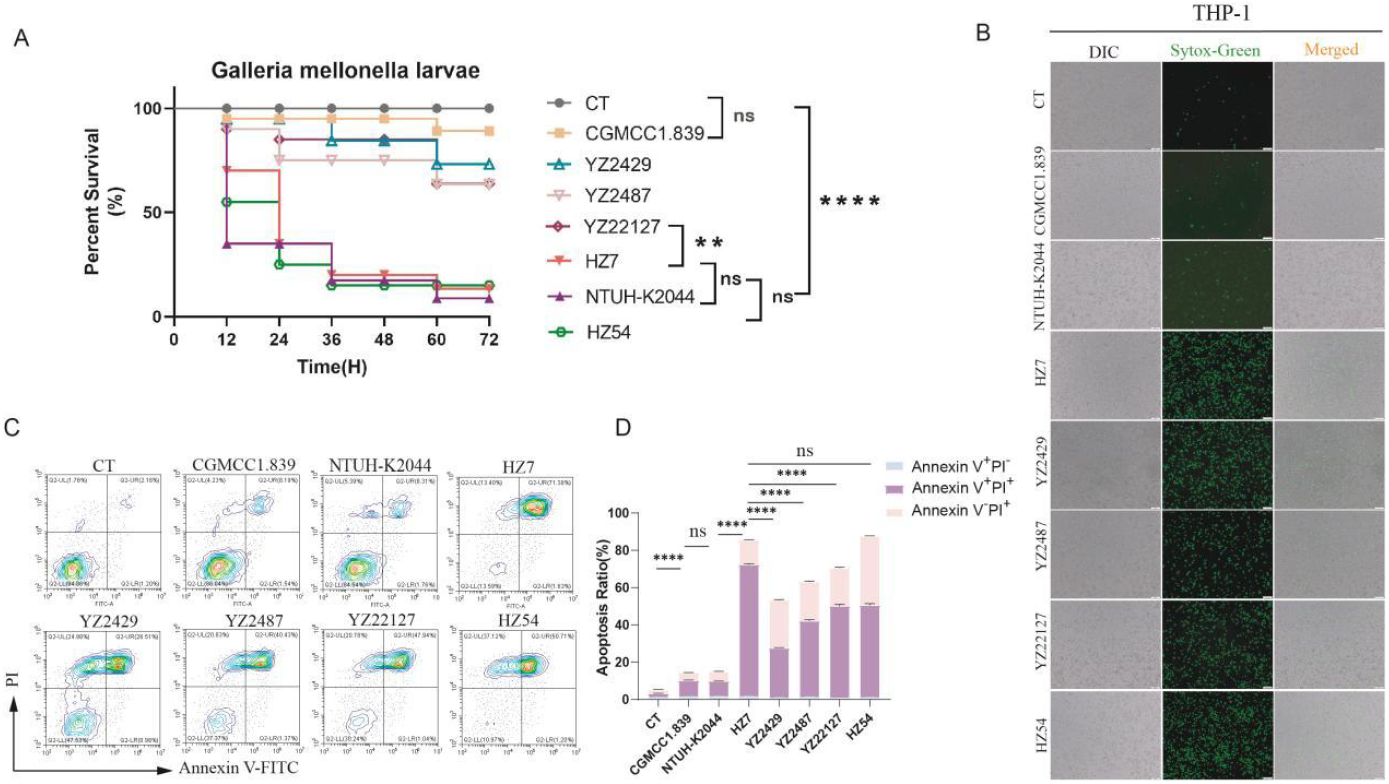
Virulence phenotype of ST617 *K. pneumoniae* strain HZ7. **(A)** The survival rates of HZ7 infected with Galleria mellonella larvae. Twenty larvae were used per strain. G. mellonella larvae were inoculated with 10 μl of each strain at doses ranging from 1×10^6^ and incubated at 30°C, and the viability of the infected larvae was assessed 72 h. Data in B is mean± SEM. ns, not significant; ****P < 0.0001, as assessed by one-way ANOVA test. **(B)** Cell death was measured by staining with SYTOX Green followed by fluorescence microscopy. THP-1 differentiated macrophages were challenged with different *K. pneumoniae* strains (MOI=10:1) at 16 hpi. Scale bars, 100 μm. **(C)** Representative flow cytometry analysis of THP-1 apoptosis (MOI=10:1) at 16 hpi. CT, uninfected.

To further evaluate the cytotoxicity of clinical isolate HZ7, we infected THP-1 differentiated macrophages and fluorescent nucleic acid stain SYTOX Green was used to assess the integrity of the plasma membranes as a measure of cell death. As shown in Figure 1(B), THP-1 infected with HZ7 and HZ54 exhibited more green fluorescence than cells infected with YZ2429, YZ2428, YZ22127, whereas very few cells were stained with SYTOX green infected with NTUH-K2044 and CGMCC at 16hpi.

In addition, we used FITC Annexin V/Dead Cell Apoptosis Kit in flow cytometry to assess cell viability of THP-1 cells infected with different strains. Consistent with the above finding, THP-1 infected with HZ7 showed the highest percentage of apoptotic cells. Interestingly, HZ7 and HZ54 showed the same proportion of dead cells, via apoptosis and necrosis (Figure 1C-D). Together, these data indicate that HZ7 was shown to be the most virulent strain among the ST617 strains, but was similar to the virulence of the HZ54 strain.

### Biofilm forming ability of *K. pneumoniae* strain HZ7

Biofilms are surface-attached groups of microbial cells, and the cells within the biofilm are encased in an extracellular matrix composed by polysaccharide, proteins and DNA[24].The ability to form biofilms can be an important virulence characteristic of *K. pneumoniae*[25]. To evaluate the biofilm formation ability of different *K. pneumoniae* strains, crystal violet was used to stain biofilm materials from different strains. The results suggest that HZ7 had the strongest biofilm formation ability, while YZ2429, YZ2487, and YZ22127 performed almost uniformly and were all weaker than HZ7 (Figure 2).

**Figure 2.**
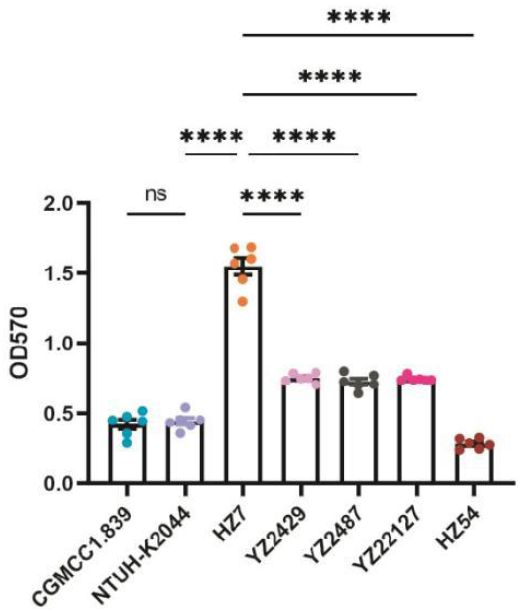
Crystal violet biofilm assay. Crystal violet (1%) was used to stain biofilm materials deposited by different *K. pneumoniae* strains. The absorbance at 570 nm of each sample was recorded. Value is indicated as means±SEM. ns, not significant; ****P < 0.0001, as assessed by one-way ANOVA test.

### Highly virulent *K. pneumoniae* strain HZ7 survived well in THP-1 cells while environmental *K. pneumoniae* strain CGCC did not

To assess intracellular survival, THP-1 differentiated macrophages were infected with the *K. pneumoniae* HZ7 strain and the control strain CGMCC1.839 and the number of surviving intracellular bacteria was quantified at different time points. As shown in Figure 3A, although bacterial numbers declined over time, the infected THP-1 macrophages were unable to completely eliminate HZ7 during the 24 h infection. In contrast, the CGMCC-infected THP-1 cells almost completely eliminated the bacteria by 4 hours. At the same time, we transformed the broad-host range fluorescence plasmid pMF230 into HZ7 and CGMCC1.839 strains, and constructed GFP-HZ7 and GFP-CGMCC1.839 strains that continuously expressed GFP fluorescent protein for easier monitoring. Thus, intracellular bacteria were visible in infected THP-1 cells (green, globular shaped). The results showed that a large amount of green fluorescence was observed in HZ7-infected THP-1 cells under fluorescence microscopy at 16 hpi, whereas green fluorescence was not found in CGMCC1.839-infected THP-1 cells (Figure 3B), indicating good survival of strain HZ7 but poor survival of the environmental strain CGMCC1.839. Taken together, these data indicate that HZ7 strain could evade immune clearance and survive inside host cells, while the environmental strain CGMCC1.839 could not.

**Figure 3.**
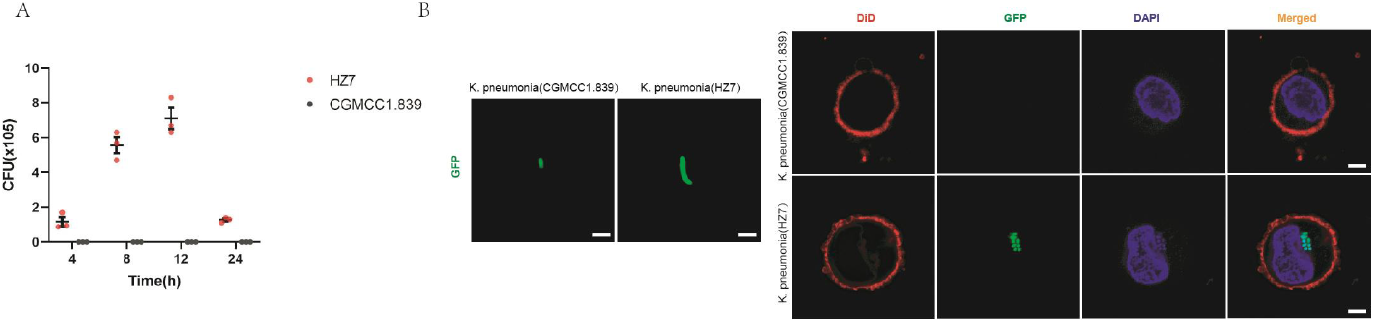
Intracellular survival of *K. pneumoniae* HZ7. **(A)** The number of CFUs in THP-1 infected cells was determined by plating samples on LB agar. The initial multiplicity of infection (MOI) was 2:1 with 3.2 × 10^6^ THP-1 cells per well. Value is indicated as means ± SEM, as assessed by unpaired two-tailed Student’s *t*-test. **(B)** Fluorescence microscopy of THP-1 cells infected with *K. pneumoniae* HZ7 and a non-pathogenic strain *K. pneumoniae* CGMCC1.839 (MOI=10:1) at 16 hpi. *K. pneumoniae* was transformed by the plasmid pMF230 with GFP green fluorescence. Cell membranes can be stained by DID solid (1,1’-Dioctadecyl-3,3,3’,3’-Tetramethylindodicarbocyanine,4-Chlorobenzenesulfonate). DAPI was used to stain the nucleus. Scale bars, 5 μm.

### Evolutionary origin of HZ7

To delve deeper into the evolutionary history of HZ7 and identify its potential origin, we retrieved all *K. pneumoniae* genomes from the NCBI database [https://www.ncbi.nlm.nih.gov/refseq]. Out of these, only 14 strains were identified as serotype ST617. Utilizing the maximum likelihood method, we constructed a phylogenetic tree based on the core genomic SNPs, with CGMCC1.839 of ST1265 serving as an outgroup. As shown in Figure 4, we found that ST617 can be divided into two distinct clades. Notably, clade 2, to which HZ7 belongs, primarily consists of strains from animal or environmental sources, while clade 1 is predominantly of human origin. This observation suggests that the origin of HZ7 might be derived from environmental or animal sources.

**Figure 4.**
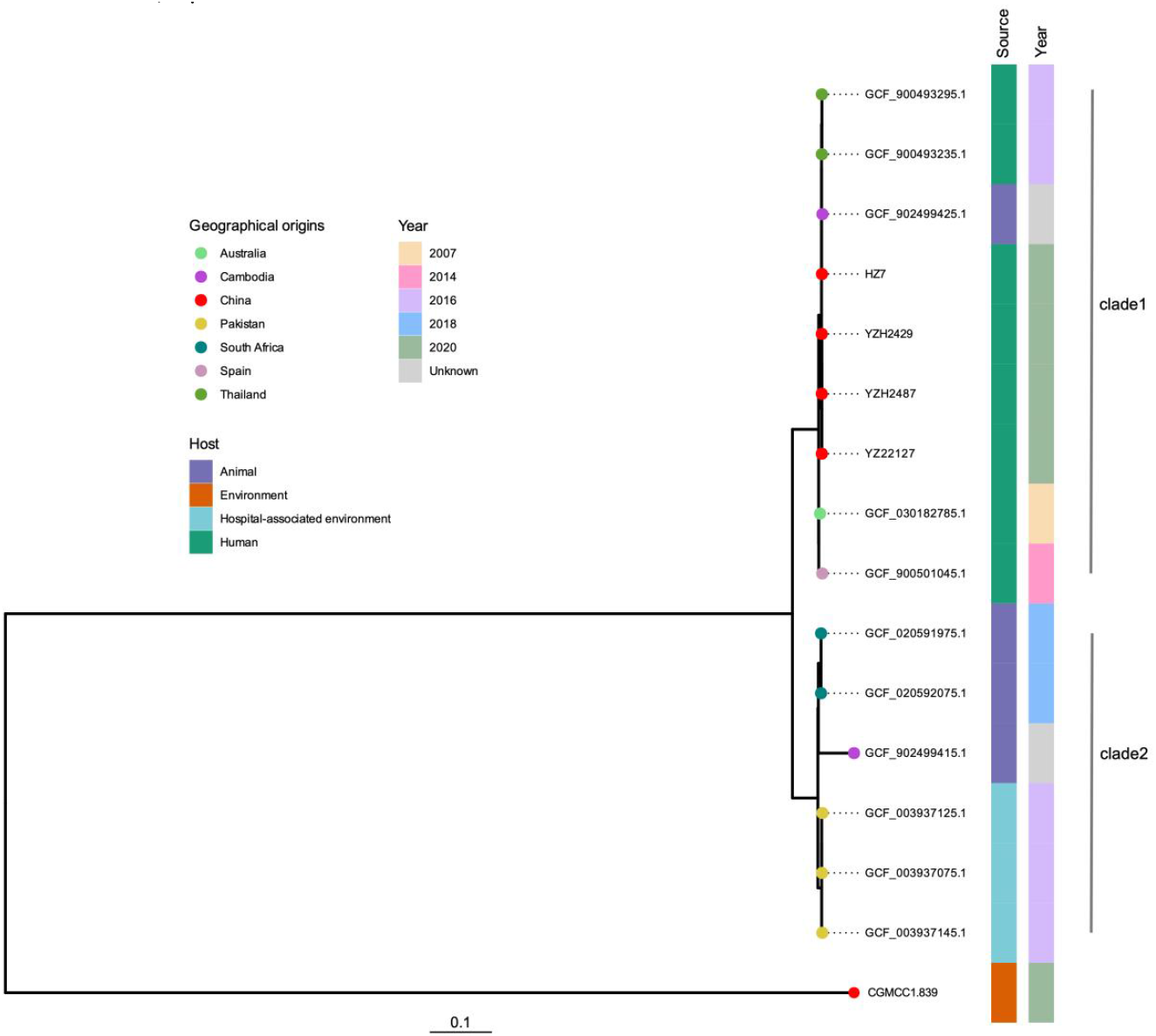
Maximum-likelihood phylogenetic analysis of ST617 isolates using core SNPs, with CGMCC1.839 serving as the outgroup. The terminal node colors correspond to the geographic origins of the respective isolates.

### Possible enhanced virulence factors in *K. pneumoniae* strain HZ7

While phylogenetic analyses suggest that *K. pneumoniae* HZ7 and the three *K. pneumoniae* YZ strains are closely evolutionarily related and possess the same virulence profile (Figure 5), our experimental data indicate a heightened virulence for strain HZ7 compared to the YZ strains derived from patient blood infections. Guided by this discrepancy, we formulated a gene presence-absence matrix with the avirulent environmetnal strain CGMCC1.839, three YZ strains, and HZ7. We particularly focused on genes unique to HZ7, considering their potential role in enhanced virulence phenotype. We pinpointed seven such distinctive genes (Table S2), which encompassed a cluster of four genes (Figure 6). This cluster is characterized by one S-type pyocin domain-containing protein and three DUF6392 family proteins. A comparative genomic analysis revealed that the quartet of ST617 strains shares an identical genetic context both upstream and downstream of this gene cluster. Notably, we identified both direct (DRs) and inverted repeat sequences (IRs) in the upstream and downstream regions of the gene cluster (Figure 6). This observation suggests a potential horizontal gene transfer event, bringing this gene cluster into the ST617 backbone in strain HZ7. This integration might have served as a catalyst for the enhanced virulence observed in HZ7.

**Figure 5.**
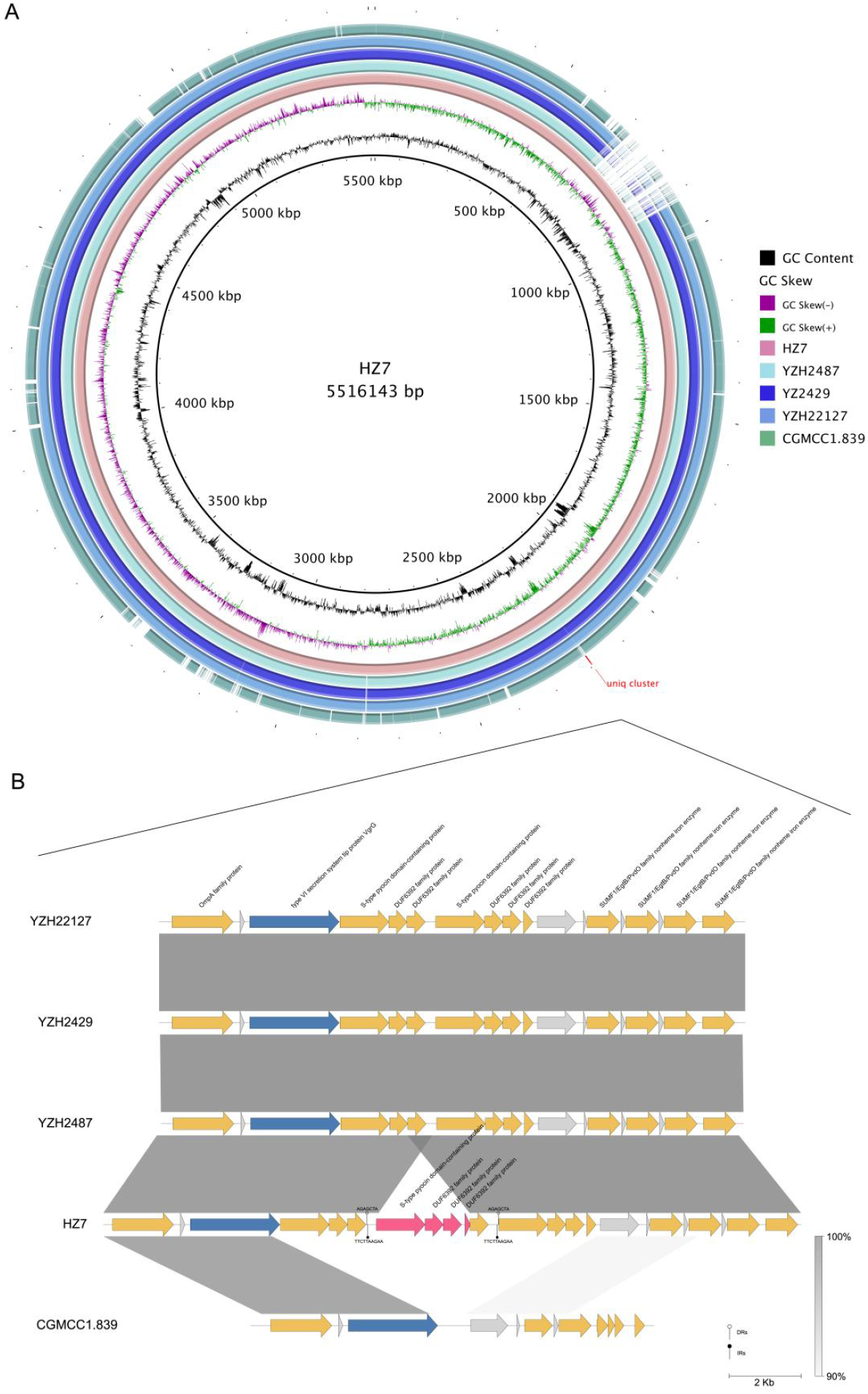
Virulence gene distribution heatmap. Colors denote the presence of genes, while blank indicates their absence. Virulence genes are categorized by function, with distinct colors representing each category.

**Figure 6.**
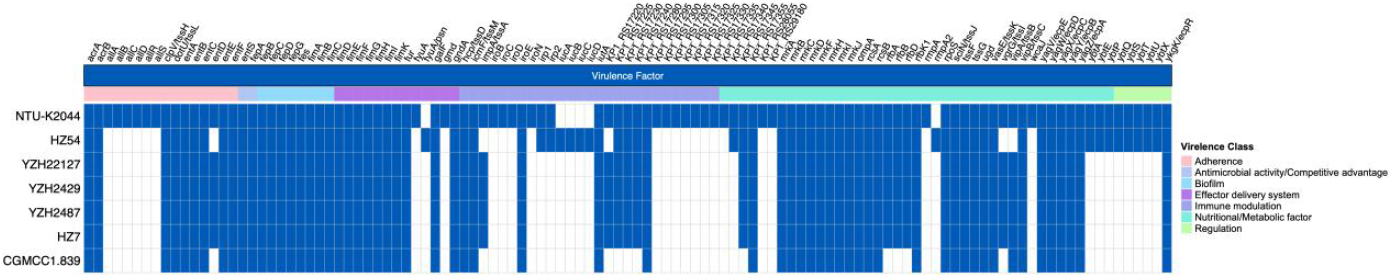
Chromosomal comparison between HZ7 and select low virulence strains. **(A)** Depicts chromosomal variances between HZ7 and strains YZ2429, YZ2487, YZ22127, and CGMCC1.839. Each concentric circle delineates the chromosomes of distinct strains; areas devoid of color signify disparities. Gene clusters exclusive to HZ7 are highlighted in red. **(B)** Illustrates the genetic landscape of distinctive gene clusters. Varying hues denote diverse gene functionalities. The gray hue indicates hypothetical proteins, while red underscores the unique gene clusters.

**Figure 7.**
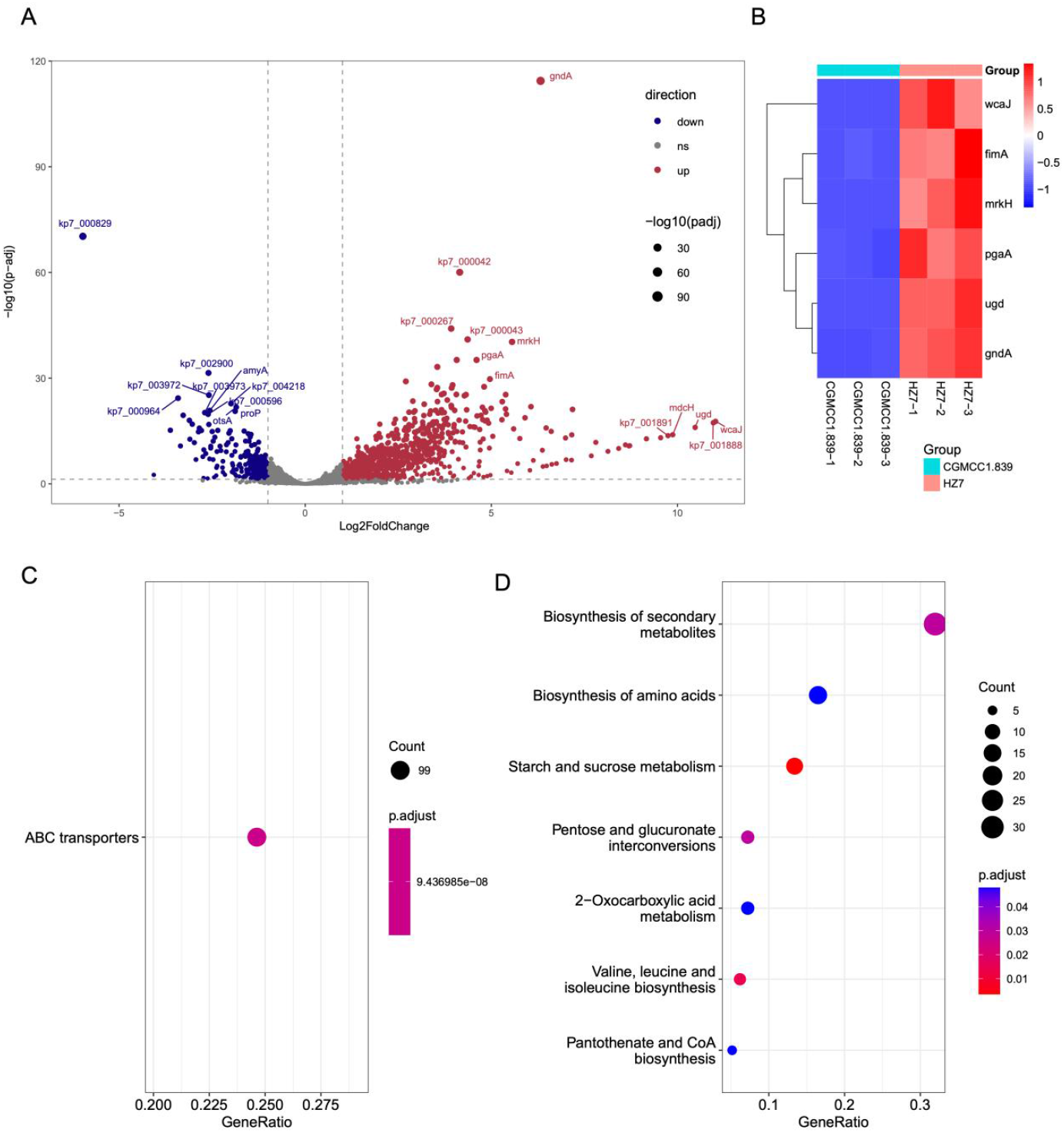
Differentially expressed genes between strong biofilm producer HZ7 and a weak biofilm producer CGMCC1.839. **(A)** Volcano plot of differentially expressed genes between *K. pneumoniae* HZ7 and CGMCC1.839. **(B)** The heat maps showing the marker gene and *wcaJ* expression. **(C-D)** Point plot of KEGG pathways for up-regulated genes.

### Transcriptome analysis of strong biofilm producer HZ7

To investigate DEGs in the strong biofilm producer HZ7, we conducted RNA-seq analysis on K. *pneumoniae* strain HZ7 and the weak biofilm producer strain CGMCC1.839. A total of 516 DEGs were identified, with 470 genes up-regulated and 47 genes down-regulated (p.adjust < 0.05, absolute log2 fold change >1) (Fig. 5A). Among the up-regulated genes, those potentially associated with biofilm formation were selected for cluster heat map analysis (Fig. 5B, Table S3). The most significantly up-regulated gene, *wcaJ*, exhibited a log2 fold change of 10.96. WcaJ, a glucosyltransferase involved in capsular polysaccharide synthesis, may play a key role in biofilm formation and contribute to the high virulence of HZ7 [26, 27]. Additionally, *gndA*, which encodes NADP-dependent phosphogluconate dehydrogenase and is located within the capsular polysaccharide synthesis (CPS) locus, has been implicated in biofilm formation [28, 29]. The deletion of *ugd*, encoding UDP-glucose 6-dehydrogenase, was found to reduce capsular polysaccharide production and decrease biofilm formation [30]. Interestingly, these DEGs were enriched in the signaling pathways of starch and sucrose metabolism and ABC transporters, both of which are involved in capsular polysaccharide synthesis known to be involved in virulence [31] (Fig. 5C, 5D). These findings suggest that the robust biofilm formation and high virulence of HZ7 are closely linked to its enhanced capsular polysaccharide synthesis.

## Conclusions

In this study, we identified a new hv-KP strain of ST617 ESBL^+^ -HZ7 that causes sepsis and showed strong virulence in various models. Compared with other reported ST617 carbapenem and tigacycline resistant strains (YZ2429, YZ2487, YZ22127)[21], HZ7 was more virulent. Meanwhile, compared with the well known hvKP strain—NTUH-K2044 and the clinically highly virulent isolate ST11-K64—HZ54, HZ7 exhibits higher virulence in cytotoxicity test and in the Galleria mellonella model, In addition, HZ7 showed strong biofilm forming ability and survived well intracellularly while the environmental strain CGMCC1.839 did not. Whole-genome sequencing and transcriptome sequencing analyses to our surprise, identified no known common hypervirulence genes in HZ7 genome. Interestingly, compared with isolates YZ2429,YZ2487,YZ22127 and non-pathogenic environmental strain CGMCC1.839, HZ7 was found to contain seven unique genes—kp7_000748, kp7_002156, kp7_002157, kp7_002158, kp7_002159, kp7_003103, kp7_003104that may cause high virulence. Transcriptome analysis of HZ7 identified six genes—*wcaJ, fimA, mrkH, pgaA, ugd*, and *gndA* that may be involved in biofilm formation. Future studies need to determine the prevalence of the hvKP ST617 ESBL+ strain and address the detailed molecular mechanism of the high virulence of this strain.

## Materials and Methods

### Bacterial strains

From 2021-2022, 100 non-repetitive *K. pneumoniae* strains from patients of the Department of Infectious Diseases, The First Affiliated Hospital of Zhejiang University were collected. All isolates were re-identified as *K. pneumoniae* using the matrix-assisted laser desorption/ionization-time of flight mass spectrometry (MALDI-TOF MS). The *K. pneumoniae* HZ7 strain was isolated from the bile duct infection of a 35-year-old male. HZ54 was isolated from the chest drainage fluid of a 63-year-old man. YZ2429, YZ2487 and YZ22127 were from the Jiangsu Key Laboratory of Zoonosis/Jiangsu Co-Innovation Center for Prevention and Control of Important Animal Infectious Diseases and Zoonoses. *K. pneumoniae* strain NTUH-K2044 was selected as a typical representative of hvKP. CGMCC1.839, which was isolated from Donghu Farm in Wuhan, was purchased from China General Microbiological Culture Collection (CGMCC).

### Antibiotic susceptibility testing

Twenty antimicrobial agents including Extended-spectrumβ-lactamase(ESBL), Imipenem, Ertapenem, Cotrimoxazole, Ceftazidime, Cefepime, Cefuroxime axetil, Cefuroxime sodium, Cefazolin, Ceftriaxone, Cefoxitin, Levofloxacin, Gentamicin, Tigacycline, Aztreonam, Ampicillin, Ciprofloxacin, Meropenem, Amikacin, Kanamycin were used for antibiotic susceptibility testing. Breakpoint determination is based on the Clinical and Laboratory Standards Institute (CLSI). *Escherichia coli* ATCC25922 was used as a quality control strain.

### Construction of HZ7.GFP strain

Broad host range GFP plasmid pMF230-Amp purchased from Beijing Biosea Biotechnology. MIC results showed that HZ7 was resistant to Ampicillin. Both HZ7 and CGMCC1.839 were sensitive to kanamycin. One-step cloning was used to modify the resistance of pMF230. PBAV1K-Kan provides the kanamycin gene template. pMF230.vec_rv: AGAGTTTGTAGAAACGCAAAAAGGCC, kan_fw:CCTTTTTGCGTTTCTACAAACTCTcaaattctatcataattgtgg, kan_rv: ctaaaacaattcatccagtaa, pMF230.vec_fw: ttactggatgaattgttttagCTGTCAGACCAAGTTTACTCAT, oriT_fw:GTGCGAATAAGGGACAGTGAA, oriT_rv: TTCACTGTCCCTTATTCGCAC. pMF230-Kan plasmid was electroporated into strains HZ7 and CGMCC1.839 (voltage 2.5kV).

### Survival rates of Galleria mellonella larvae infected with *K. pneumoniae* strains

*K. pneumoniae* strains were cultured in LB medium with shaking at 200 rpm for 16h at 37°C. Bacterial suspensions were collected with PBS. 10ul was injected into the left hind proleg of the larvae with a 25 μl microinjector, approximately 10^6^ colony-forming units (CFU) per group of strains. Each group had 20 larvae, which were then placed on a 150mm petri dish containing filter paper and cultured in the dark at 37°C. Larvae survival was observed and recorded every 12 hours. The survival curve was plotted using GraphPad Prism 9.3.0.

### Annexin V-FITC/PI staining

The human monocytic cell line THP-1 cells were seeded into 12-well plates with a cell density of 5 × 10^5^ cells/ml. THP-1 were differentiated to macrophage-like characteristics cells by phorbol 12-myristate 13-acetate (PMA,1μM). The MOI of bacterial infection to THP-1 cells was 10:1. Cells were collected 16h after infection and treated with Annexin V-PI apoptosis detection kit(MULTI SCIENCES, AT101-100). Flow cytometry detection was performed using Beckman CytoFlex instruments and data analysis using CytExpert software.

### SYTOX Green nucleic acid staining

SYTOX Green Nucleic Acid Stain is a nucleic acid dye with high affinity, which can easily penetrate into the damaged cytoplasm membrane but cannot penetrate the living cell membrane. THP-1 cells(5 × 10^5^ cells/ml) were infected with different strains of *K. pneumoniae* at MOI= 10:1 for 16h. The dead cells were then stained with SYTOX Green nucleic acid (167 nM). Fluorescence microscopy was used for detection (Excitation/Emission is 504/523).

### Intracellular survival assay

THP-1 cells were placed on a 6-well plate with a cell density of 8 × 10^5^/ml. Treatment with Phorbol 12-myristate 13-acetate (PMA) for 24h induced cell adhesion. 2h after infection (MOI= 2:1), the cells were washed twice with gentamicin containing 100 μg/ml. Cells were collected after incubation with 100 μg/ml of gentamicin containing 10% FBS for 4h, 8h, 12h and 24h. Cells were lysed with 1% Triton-100 for 5 min and CFU counting was performed.

### Flow cytometry

In order to detect the survival of *K. pneumoniae* HZ7 strain in host cell, we electroporated pMF230-Kan.GFP plasmid into HZ7 and CGMCC1.839 strain to make them stably express GFP fluorescence. THP-1 cells were infected with GFP plasmid containing HZ7 and CGMCC1.839 (MOI= 10:1) for 16h and then fixed with 4% Paraformaldehyde (PFA). Cell membranes were stained by DID (1, 1 ‘-Dioctadecyl - 3 filling’, 3 ‘-Tetramethylindodicarbocyanine, 4 - Chlorobenzenesulfonate Salt, 10µM), and DAPI was used to stain the nucleus. Fluorescence detection was performed using a Zeiss 900 inverted laser confocal microscope.

### Biofilm formation

The ability of biofilm formation was detected by crystal violet staining. The bacteria were diluted 100 times with LB broth and inoculated into 96-well plates. After incubation at 37 ° C for 48h, the planktonic bacteria were washed with phosphate buffered saline (PBS). After the removal of PBS, the 96-well plate was fixed in an oven at 60°C for 1h, stained with Crystal violet (1%w/v) for 15min, washed with PBS for 3 times, followed by dissolving crystal violet with 95% ethanol. The absorbance at 570 nm of each sample was recorded. Six replicates were used per group.

### Whole-genome sequencing, assembly, annotation, and bioinformatic analysis

Genomic DNA was procured utilizing the AxyPrep bacterial genomic DNA miniprep kit (Axygen Scientific, Union City, CA, USA). The complete genomes of HZ7 and CGMCC1.839 were constructed by amalgamating short-read information derived from Illumina sequencing with the long-read data procured from Oxford Nanopore sequencing via the Unicycler hybrid assembly pipeline[32]. Annotation of all genomes was conducted using the NCBI Prokaryotic Genome Annotation Pipeline (PGAP)[33]. Unique genes related to HZ7 were scrutinized employing Panaroo v1.3.2[34]with the default parameters. Core Single nucleotide polymorphisms (SNPs) across all ST617 genomes were delineated using Snippy v1.14.6 (https://github.com/tseemann/snippy), designating CGMCC1.839 as the outgroup and *K. pneumoniae* HZ7 as the referential genome. A maximum likelihood (ML) phylogeny was constructed based on the core-genome alignment, utilizing RaxML v8.2.12[35]with the GTR nucleotide substitution model accompanied by 1,000 rapid bootstrap iterations. The resultant phylogenetic dendrogram was represented via the ggtree[36]. The genetic contextual analysis was compared through MUMmer[37]and rendered visually using pyGenomeViz (https://github.com/moshi4/pyGenomeViz). Comparative genomic circular maps of *K. pneumoniae* HZ7 were generated via the BLAST Ring Image Generator (BRIG) tool[38]. Virulence genes were predicted using ABRicate (https://github.com/tseemann/abricate) and visualized using the ComplexHeatmap package[39].

### Transcriptome and data analysis

Total RNA was extracted from one biofilm-proficient strain, HZ7, and one weak biofilm-producing strain, CGMCC1.839, using the Bacteria Total RNA Isolation Kit, following the manufacturer’s protocol. Three replicates were prepared for each strain. Sequencing was conducted on an Illumina NovaSeq platform, generating 150 bp paired-end reads. After quality trimming of raw reads using fastp, the reads were aligned to the HZ7 reference genome. Gene-level quantification was performed using featureCounts, and differentially expressed genes (DEGs) were identified with DESeq2, applying thresholds of an absolute log2 fold change >1 and a Benjamini–Hochberg adjusted P-value (p.adjust) < 0.05. DEGs were visualized using a volcano plot generated with the R package ggrepel. Functional enrichment analysis of up-and down-regulated DEGs was carried out using clusterProfile[40], focusing on Kyoto Encyclopedia of Genes and Genomes (KEGG) pathways.

## Acknowledgements

This study was supported by National Infectious Disease Medical Center startup fund (B2022011-1), and Jinan Microecological Biomedicine Shandong Laboratory project (JNL-2022050B).

## Author contributions

Yuyun Yu, Conceptualization, Methodology, Investigation and Formal Analysis, Writing – original draft | Xu Dong, Data curation, Investigation, Software, Visualization | Yanghui Xiang, Methodology, Validation | Yi Li, Methodology, Validation | Kefan Bi, Software | Jiaying Liu, Methodology | Lin Sun, Resources | Tiantian Wu, Investigation | Ying Zhang, Conceptualization, Writing – review and editing

